# No effect of hippocampal lesions on stimulus-response bindings

**DOI:** 10.1101/111898

**Authors:** Richard N. Henson, Aidan J. Horner, Andrea Greve, Elisa Cooper, Mariella Gregori, Jon S. Simons, Sharon Erzinçioğlu, Georgina Browne, Narinder Kapur

## Abstract

The hippocampus is believed to be important for rapid learning of arbitrary stimulus-response contingencies, or S-R bindings. In support of this, Schnyer *et al.*, 2006 (Experiment 2) measured priming of reaction times (RTs) to categorise visual objects, and found that patients with medial temporal lobe damage, unlike healthy controls, failed to show evidence of reduced priming when response contingencies were reversed between initial and repeated categorisation of objects (a signature of S-R bindings). We ran a similar though extended object classification task on 6 patients who appear to have selective hippocampal lesions, together with 24 age-matched controls. Unlike Schnyer *et al.*, 2006, we found that reversing response contingencies abolished priming in both controls and patients. Bayes Factors provided no reason to believe that response reversal had less effect on patients than controls. We therefore conclude that it is unlikely that the hippocampus is needed for S-R bindings.

## Introduction

The medial temporal lobes (MTL), and hippocampus in particular, are thought necessary for rapid acquisition of new associations (Squire, 1992; Cohen & Eichenbaum, 1994; Schacter and Tulving, 1994; Giovanello, Schnyer, & Verfaellie, 2004). On the other hand, such MTL regions do not appear necessary for all types of rapid plasticity, such as that presumed to underlie implicit memory phenomena like priming, which can also occur after a single exposure to a stimulus (e.g., Cave and Squire, 1992; Schacter *et al.*, 1993). Priming is often measured by decreases in the reaction time (RT) to perform a simple classification task on a stimulus, such as deciding whether the object depicted by a picture is large or small in real life. Such RT priming has often been associated with facilitated perceptual or conceptual processing, occurring in cortical regions outside the MTL (Henson, 2003).

However, recent studies have shown that the dominant cause of such classification-based RT priming is the encoding and retrieval of Stimulus-Response (S-R) bindings (see Henson *et al.*, 2014, for a recent review). According to this account, the response made to the first presentation of a stimulus is bound together with that stimulus, such that when that stimulus is repeated, the response can be retrieved. This retrieval of a previous response is assumed to be faster than repeating the original perceptual/conceptual processing that generated the response on the initial stimulus presentation, causing the RT priming. However, if the task changes between initial and repeated presentations, such that the response is changed, the amount of RT priming is reduced (relative to “novel”, control stimuli that were not presented in the original task). Indeed, sometimes priming is abolished by a response reversal, or even becomes negative priming, i.e, slower RTs for repeated than novel stimuli, possibly owing to interference from retrieval of incorrect responses (Horner and Henson, 2011).

Neuroimaging data support the contribution of rapidly learnt S-R bindings to performance on classification tasks. Several fMRI studies in healthy individuals have found that the decreased fMRI response following repetition of visual stimuli (“repetition suppression”, RS), which has been associated with priming (Koutstaal *et al.*, 2001; Schacter and Buckner, 1998; Simons *et al.*, 2003), is reduced when the classification task is reversed. This reduction in RS following response reversal has been seen in lateral prefrontal regions commonly associated with response selection, and occasionally in ventral temporal regions commonly associated with perceptual/conceptual component processes (Dobbins *et al.*, 2004; Horner and Henson, 2008; Race *et al.*, 2009), though is not readily apparent in MTL regions.

Given that a typical priming experiment entails tens if not hundreds of unique stimuli, the retrieval of the appropriate S-R binding when one of those stimuli is repeated suggests that the brain has an impressive capacity to store many such S-R bindings. To test whether this capacity for rapid learning of multiple, unique S-R associations is supported by MTL, Schnyer *et al.* (2006; Experiment 2) reported priming data from a speeded classification task on a group of 9 patients with MTL damage, together with age-matched controls. Participants were initially asked to decide “Is the object bigger than a shoebox?”, but then after one or three presentations of each stimulus, the task reversed to “Is the object smaller than a shoebox?”. Controls showed the usual reduction in RT priming when the task was reversed, indicative of S-R bindings. RT priming in the patients however showed no detectable effect of the task being reversed (see ahead to Figure 3 for a re-plotting of Schnyer *et al’s* data). The authors therefore concluded that MTL regions are responsible for S-R learning.

Though MTL damage was “radiologically-verified” in each patient, the extent of that damage was not reported by Schnyer *et al.* (2006), so they were unable to conclude whether S-R bindings are supported specifically by the hippocampus, or by other MTL regions like entorhinal, perirhinal or parahippocampal cortices. We recently reported six patients whose MRI scans showed clear evidence of hippocampal volume reduction, with little sign of gray-matter damage outside the hippocampus (Henson *et al.*, 2016). Our main aim in the present experiment was therefore to determine whether the S-R deficit reported by Schnyer *et al* is selective to hippocampal damage.

Our second aim was to test whether Schnyer *et al’s* results generalise to a modified version of the visual classification priming task, which we have previously shown to additionally identify the contribution of Stimulus-Classification (S-C) bindings. S-C bindings are not controlled in Schnyer *et al’s* task reversal paradigm (see Horner and Henson, 2011). This may be important because we have previously shown that S-R bindings include multiple levels of response representation (Horner and Henson, 2009) and multiple levels of stimulus representation (Horner and Henson, 2011, 2012).

Our alternative paradigm (initially proposed by Denkinger and Koutstaal, 2009) involves keeping the task constant (e.g, “Is the object bigger than X?”), but changing the referent instead (i.e, X). This paradigm simultaneously reverses multiple levels of response representations (Horner and Henson, 2009; see also Schnyer *et al.*, 2007; Dennis and Perfect, 2012), as illustrated in Figure 1. In this example, the response associated with an object (e.g, monkey) when it is judged to be bigger than a shoebox could include the specific motor Action (e.g, right index finger press), the Decision (e.g, “yes”/”no”) and/or the Classification label (e.g, “bigger”/”smaller”). Reversing the task, as done by Schnyer *et al,* potentially disrupts the value of retrieving the previous Action or Decision, but retrieving the previous Classification (e.g., “bigger”) could still help generate a response (e.g, “no” to the reversed task of smaller than a shoebox), by-passing the need for extensive perceptual or conceptual processing. However, changing the referent instead, for example to a wheelie bin (Figure 1), additionally disrupts the value of retrieving a prior Classification, as shown by Horner and Henson (2009).

**Figure 1.**
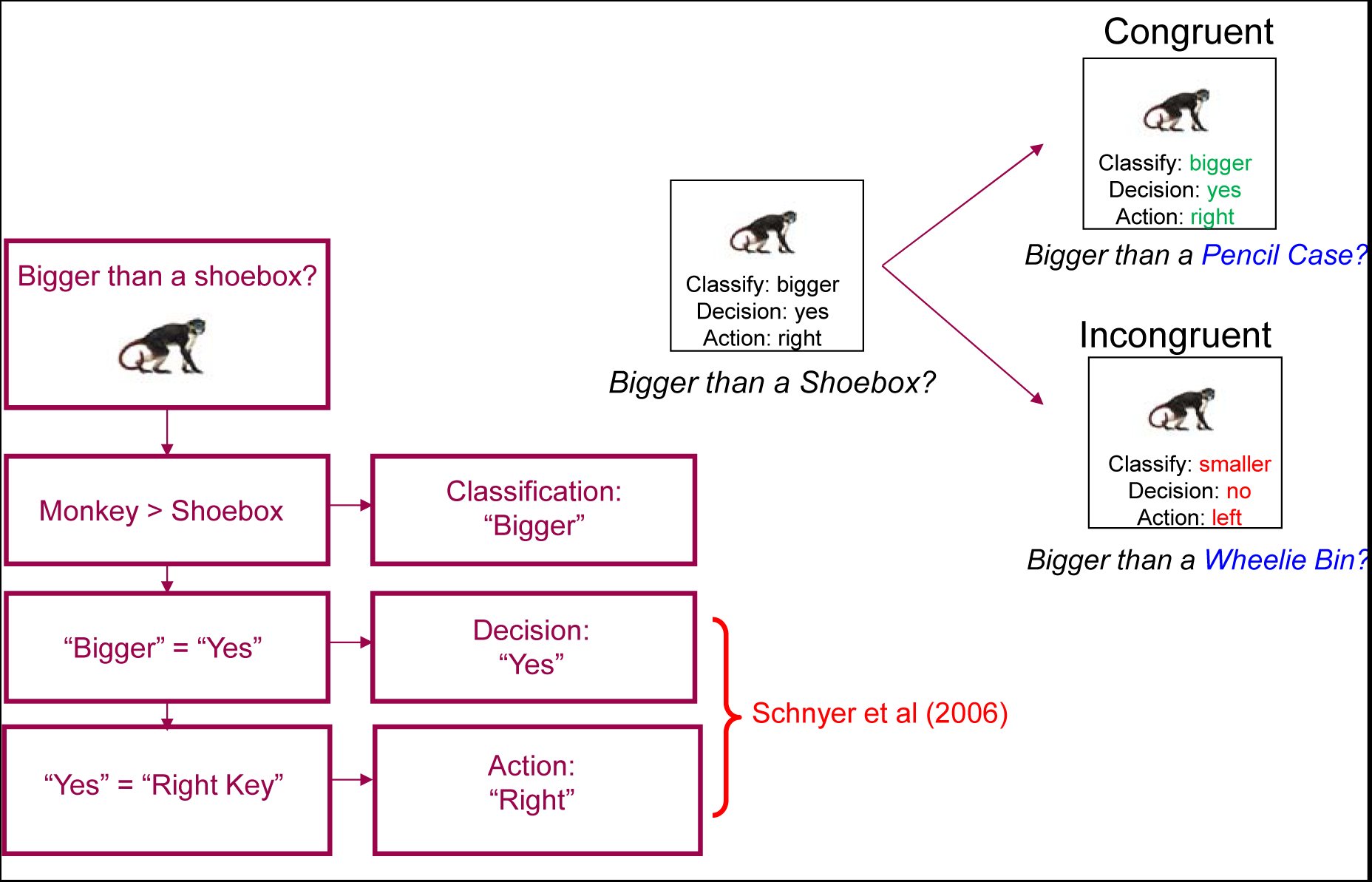
Bottom Left: Schematic of possible response representations (Classifications, Decisions and Actions) that could be bound with a stimulus in a classification task. Reversing the task, e.g., from “Bigger than a shoebox” to “Smaller than a shoebox”, as in Schnyer et al. (2006), reverses the Decision and Action, but not the Classification. Top Right: Changing the referent (e.g, from a shoebox to a wheelie bin), on the other hand, reverses all three levels of response representation.

Furthermore, we can also test the type of stimulus representation in S-R bindings by orthogonally varying whether or not the stimulus is repeated in the same perceptual form (e.g, picture or name) as its initial presentation. We previously showed evidence for two levels of stimulus representation: a form-specific and more abstract representation (Horner and Henson, 2011; see also Allenmark *et al.*, 2015; though see Schnyer *et al.*, 2007). We included this “Within-format” versus “Across-format” manipulation in the present experiment to test whether patients are similarly able to form S-R bindings that abstract away from the precise stimulus form. Indeed, the present experiment is identical to that in Experiment 1 of Horner and Henson (2011), except that: 1) we tested older healthy controls and patients, rather than young controls, 2) the trials were self-paced rather than running at a fixed rate, to make the task easier for patients (and older controls), who generally respond slower and show greater variability, and 3) used two rather than three presentations of each stimulus before the referent change, to try to maintain the same total duration as our previous experiment.

More precisely, Experiment 1 conformed to a 2x3x2 factorial design, with between-subject factor Group (N=24 Controls vs N=6 Patients) and within-subject factors: Study Condition (Within-format Primed, Across-format Primed, Novel) and Congruency (Congruent, Incongruent; see Methods section for how the Novel condition was split into Congruent and Incongruent conditions). Like Horner and Henson (2011), we analysed priming in both subtractive (Novel – Primed) and proportional ((Novel – Primed) / Novel) terms, but focus on the proportional measure because this was the measure used by Schnyer et al. (2006) to allow for the fact that patients tend to have longer overall RTs than controls. Once priming scores have been calculated, the design equates to a 2 (Group) × 2 (Format) × 2 (Congruency) factorial design. Based on Schnyer et al’s findings, we expected an interaction between Group and Congruency on the amount of priming, with Controls showing a greater effect of Congruency than Patients. More specifically, we predicted that Controls would show greater priming than Patients in Congruent trials (because Controls but not Patients benefit from S-R bindings), but comparable or even less priming than Patients for Incongruent trials (where Controls would either ignore S-R bindings, or experience interference from incompatible S-R bindings).

## Materials and Methods

### Participants

The 6 patients were selected from the Cambridge Hippocampal Panel, and are the same as those reported in Henson *et al.* (2016). The study was approved by NRES Ethics Committee East of England (ref 12/EE/0190) and written consent obtained according to the Declaration of Helsinki. The patients were referred on the basis of reported memory difficulties and, in some cases, a diagnostic MR scan that showed an indication of limited MTL damage, with various aetiologies. More detailed research MRI scanning revealed that the hippocampus was the only brain region showing consistent gray-matter loss across all 6 patients (see Henson *et al.*, 2016, for more details).

Twenty-four control participants were recruited from the Volunteer Panel of the Medical Research Council (MRC) Cognition and Brain Sciences Unit (CBU). There was no significant difference in mean age of these controls (M=60, range 50-72) and that of the patients (M=58, range 39-66), t(28)=0.54, p=.60. Thirteen of the control group were female, whereas only one patient was female, and therefore analyses below were repeated with sex as a covariate of no interest.

### Materials

Stimuli were 384 coloured images of everyday objects and their names, previously used by Horner & Henson (2009), split into two groups, relating to the wheelie bin and pencil case referent change (192 stimuli per group). For the wheelie bin referent group, stimuli were classified so that 25% were smaller than both a shoebox and a wheelie bin (Congruent), 50% were bigger than a shoebox but smaller than a wheelie bin (Incongruent) and 25% were bigger than both a shoebox and a wheelie bin (Congruent). For the pencil case referent group, 25% were smaller than a pencil case and a shoebox (Congruent), 50% were bigger than a pencil case but smaller than a shoebox (Incongruent) and 25% were bigger than a pencil case and a shoebox (Congruent). This resulted in 96 stimuli per Congruency condition for each referent group. Stimuli within each of these Congruency groups were randomly assigned to one of three Study Condition groups, relating to whether they were presented as a picture at Study (Within-format Primed), a word at Study (Across-format Primed) or were experimentally Novel (Novel). This resulted in 64 stimuli per each of the 3 conditions when collapsing across the two referent changes. The assignment of stimuli to the three Study Condition factors was rotated across control participants.

### Procedure

Prior to the experiment, participants performed a practice session using the “bigger-than-shoebox” task, where it was made clear that this comparison referred to the object’s typical size in real life. Participants responded using a “yes” or “no” key with their right or left index finger respectively, and were required to respond as quickly as possible without compromising accuracy. Stimuli in the practice session were 10 objects (5 pictures, 5 words) that were not included in the main experiment. Following the practice session, participants were shown example photos of each object referent (i.e., shoebox, wheelie bin, pencil case) and were asked to report the average size of each referent. They were told the referent may change in the course of the experiment; however were not informed as to when this might occur.

The experiment consisted of four alternating study-test cycles (two relating to the wheelie bin referent change and two relating to the pencil case referent change) with each cycle lasting approximately 15 min. During each study phase, 64 stimuli were shown two times resulting in 128 trials. 32 stimuli were presented as pictures (Within-format) and 32 were presented as words (Across-format). Words were presented in black on a white background with the same pixel dimensions as the pictures. Each set of 32 stimuli consisted of equal numbers of Congruent and Incongruent items. Apart from ensuring no immediate repetitions, the stimulus presentation order was randomized. Participants were always asked “is the object bigger than a shoebox?” at Study.

During each Test phase, the 64 stimuli from the Study phase (Within-format and Across-format) were randomly intermixed with 32 new stimuli (Novel). All items at Test were presented as pictures. Participants were either asked “is the object bigger than a wheelie bin?” or “is the object bigger than a pencil case?”. The order of task (i.e., referent change) was counterbalanced across participants in an ABBA/BAAB manner. When combined with the 3 stimulus sets, this meant 6 different counterbalancings (though owing to an experimenter error, only 5 of the 6 counterbalancings were used for the patients, with two patients having the same stimulus assignment).

Each trial consisted of a 500ms fixation cross followed by a stimulus that remained onscreen until the participant responded, followed by a blank screen for 200ms. A response was required before the next trial started (i.e, the task was self-paced).

### Analyses

Trials with RTs less than 400ms, or two or more standard deviations above or below a participant’s mean for a given task, were excluded from the RT analyses (also rendering the RT distributions more Gaussian). Given that there is some subjectivity in determining whether an object is bigger than a shoebox, wheelie bin or pencil case, errors were defined by a difference from the modal response across participants for each object in the Horner and Henson (2011) study.

Note that for Novel stimuli, “congruency” refers to whether the correct response for the “bigger”/”smaller” task would be the same or different for the study-task referent as for the test-task referent, even though participants never actually classified Novel items according to the study-task referent. Therefore the subtraction of Novel RTs from Repeated RTs for Congruent and Incongruent conditions separately means that priming effects were not confounded by item differences owing to how “close” in size each object was to the relevant referent (see Horner & Henson, 2011, for further details and analyses).

Error rates and RTs for correct trials at Test constituted the main dependent variables. Given the focus on S-R effects, RTs were further restricted to objects also given a correct judgment on every occurrence at Study. These variables were subjected to repeated-measures analyses of variance (ANOVAs). Only ANOVA effects that survived an alpha of .05 were reported, and unless stated otherwise, t-tests were two-tailed.

Two ANOVAs were performed: 1) a two-way ANOVA on Novel trials, with factors Group (Controls vs Patients) and Congruency (Congruent vs Incongruent), and 2) a threeway ANOVA on priming scores, with factors Group, Congruency and Format Match (Within vs Across format, ie whether items depicted as Pictures at Test had been seen as Pictures or Words respectively at Study). Because of potential differences in Congruent and Incongruent Novel RTs (as tested by the first ANOVA), the two-way ANOVA on RT priming scores were performed for both subtractive and proportional definitions of priming. Subtractive priming is simply the difference in RTs for Repeated vs. Novel stimuli (Novel – Repeated); proportional priming is the difference in RTs for Repeated vs. Novel stimuli divided by Novel RTs ((Novel – Repeated) / Novel), calculated for each participant. However, we focus on the proportional priming results because: 1) this was the measure used by (Schnyer *et al.*, 2006), to allow for the fact that patients tend to respond slower overall, and 2) to allow for the fact that Incongruent Novel RTs were longer than Congruent RTs. (The low numbers of errors made a proportional measure of error priming inappropriate.)

To ease comparison with Schnyer et al (2006; Table 3), but in minor departures from Horner et al (2011; Table 1), we report standard errors rather than standard deviations, and express proportional priming in terms of %.

## Results

### Errors

The percentages of errors are shown in Table 1 (note that “errors” in this subjective size judgment are defined relative to the modal response across participants; see Methods). The error rate in controls was very similar to those in Table 1 of Horner and Henson (2011), even though the present controls were considerably older. The error rate was also very similar between the present controls and the patients, suggesting that the patients could perform the task at a similar level.

**Table 1.**
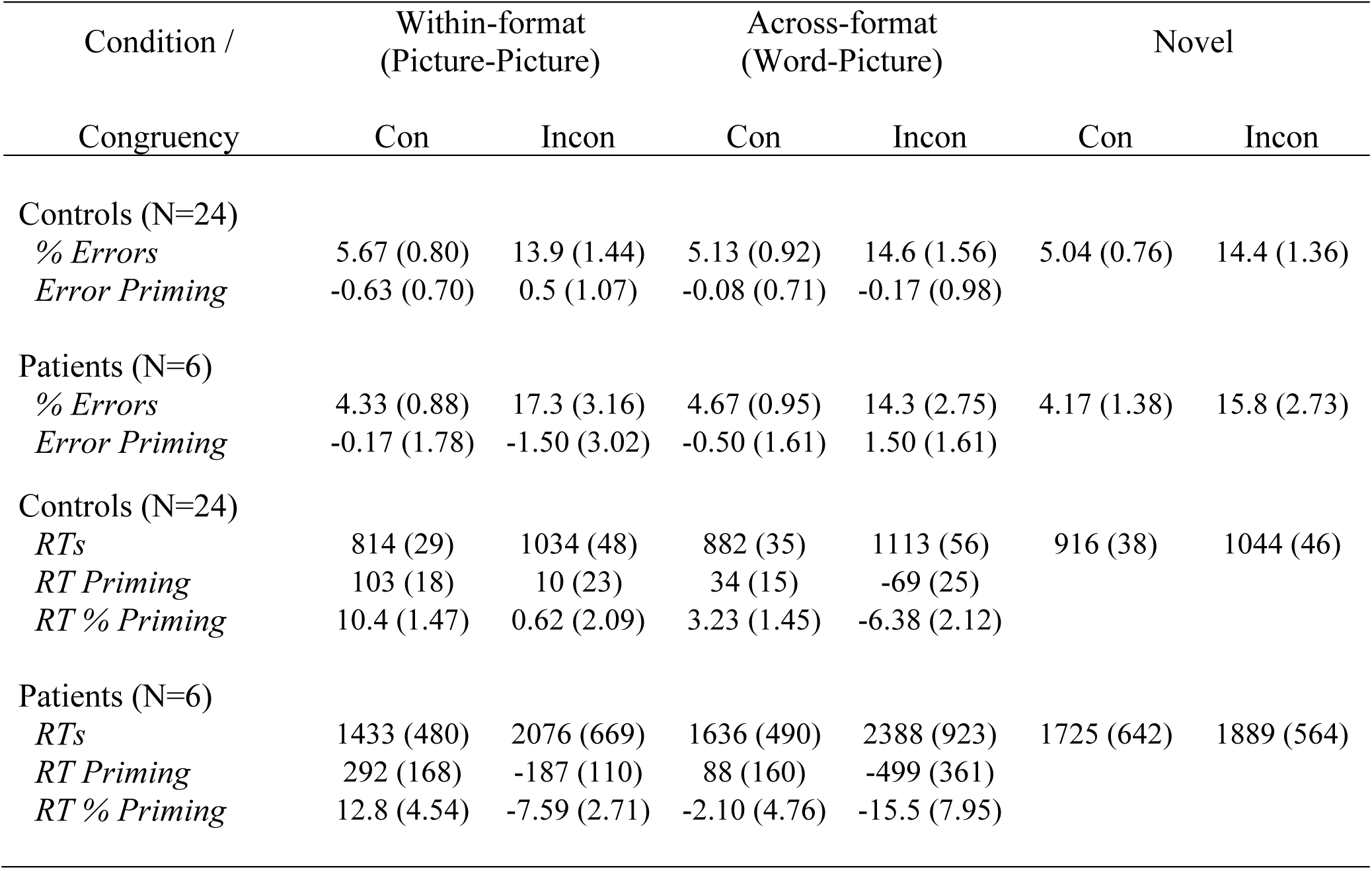
Mean percentage errors and reaction times (RTs), with standard errors in parentheses, for Within-format, Across-format and Novel conditions of Experiment 1, plus error priming, RT priming and proportional priming (% Priming) as a function of Congruency (Con, Incon). Note that for Novel stimuli, “congruency “ refers to whether the correct response for the “bigger”/”smaller” task would be the same or different for the study-task referent as for the test-task referent, even though participants never actually classified Novel items according to the study-task referent.

| Condition /<br>Congruency | Within-format<br>(Picture-Picture) |  | Across-format<br>(Word-Picture) |  | Novel |  |
| --- | --- | --- | --- | --- | --- | --- |
|  | Con | Incon | Con | Incon | Con | Incon |
| Controls (N=24) |  |  |  |  |  |  |
| % Errors | 5.67 (0.80) | 13.9 (1.44) | 5.13 (0.92) | 14.6 (1.56) | 5.04 (0.76) | 14.4 (1.36) |
| Error Priming | -0.63 (0.70) | 0.5 (1.07) | -0.08 (0.71) | -0.17 (0.98) |  |  |
| Patients (N=6) |  |  |  |  |  |  |
| % Errors | 4.33 (0.88) | 17.3 (3.16) | 4.67 (0.95) | 14.3 (2.75) | 4.17 (1.38) | 15.8 (2.73) |
| Error Priming | -0.17 (1.78) | -1.50 (3.02) | -0.50 (1.61) | 1.50 (1.61) |  |  |
| Controls (N=24) |  |  |  |  |  |  |
| *RTs* | 814 (29) | 1034 (48) | 882 (35) | 1113 (56) | 916 (38) | 1044 (46) |
| *RT Priming* | 103 (18) | 10 (23) | 34 (15) | -69 (25) |  |  |
| *RT % Priming* | 10.4 (1.47) | 0.62 (2.09) | 3.23 (1.45) | -6.38 (2.12) |  |  |
| Patients (N=6) |  |  |  |  |  |  |
| *RTs* | 1433 (480) | 2076 (669) | 1636 (490) | 2388 (923) | 1725 (642) | 1889 (564) |
| *RT Priming* | 292 (168) | -187 (110) | 88 (160) | -499 (361) |  |  |
| *RT % Priming* | 12.8 (4.54) | -7.59 (2.71) | -2.10 (4.76) | -15.5 (7.95) |  |  |

The 2×2 ANOVA on error rates in Novel conditions showed only a significant main effect of Congruency, F(1,28) = 41.7, p<.001, which reflected more errors in the Incongruent than Congruent condition. This was as expected, since these items tended to be closer to the referents and hence more ambiguous, and illustrates the importance of having separate Novel baselines with which to measure priming. There was no significant effect of Group (Controls vs Patients), nor interaction between Group and Congruency, F(1,28)’s<1.

The 2×2×2 ANOVA on subtractive priming in error rates showed no significant effects or interactions between Format Match, Congruency and Group, F(1,28)’s < 2.88, p’s > .10. These results suggest that the RT priming effects below are unlikely to reflect a speed-accuracy trade-off.

The results of the same analyses repeated with sex as a covariate were very similar (see Appendix).

### Reaction Times

In additional to the errors described above, we excluded another 8% of trials in the Control group and 2% in the Patient group with outlying RTs in Test trials or inconsistent responses across Study trials (see Methods). As expected, the present controls were slower (Table 1) than their younger counterparts in Table 1 of Horner and Henson (2011), which might explain why their subtractive priming scores were slightly larger, though their proportional priming scores were more comparable. Patients were slower still (than the present controls), though there was large variability across patients (see Appendix Table 1 for scores for each patient). Nonetheless, proportional priming was similar in size to the controls, if slightly more extreme (Figure 2).

**Figure 2.**
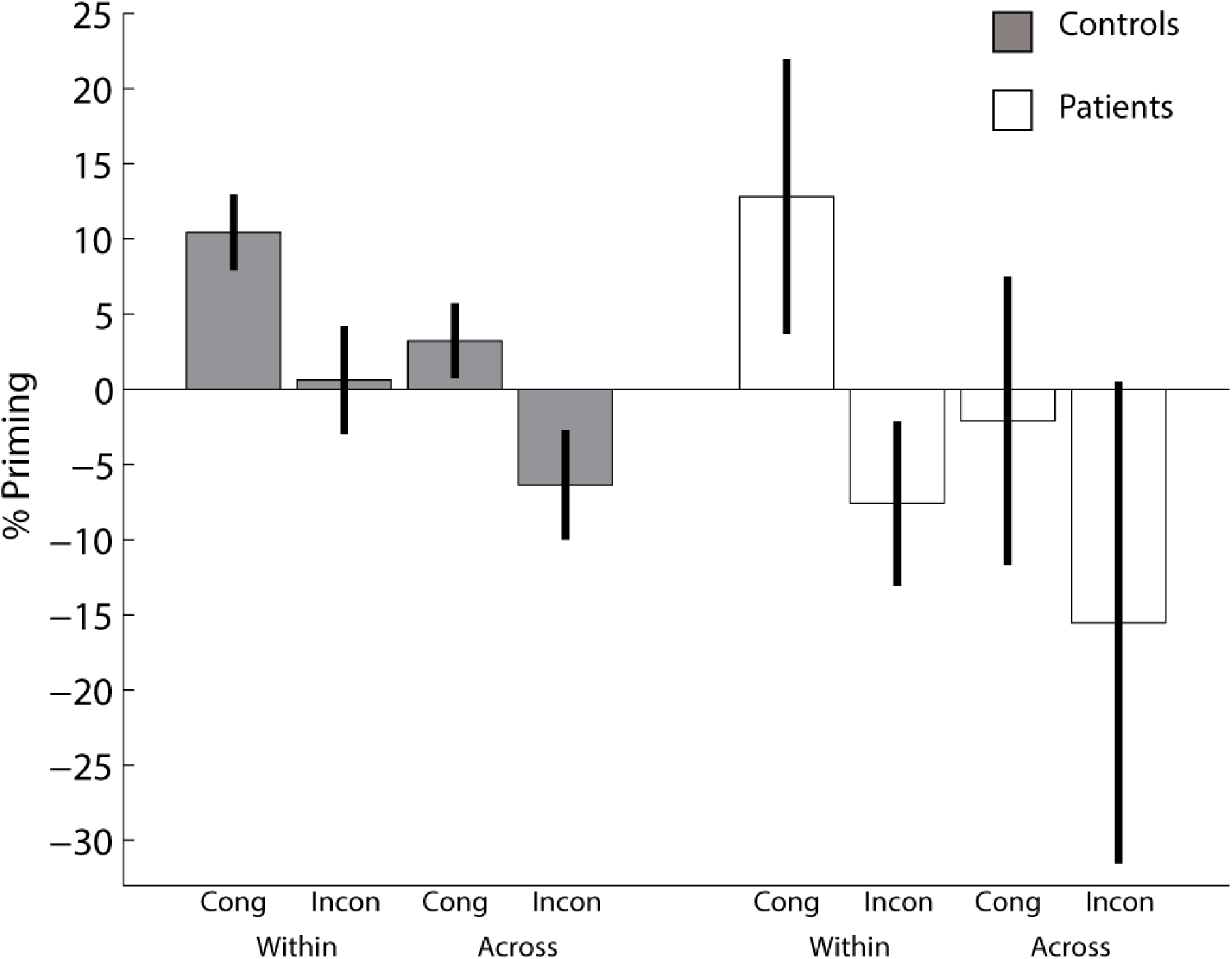
Proportional Priming for each condition and group. Cong=Congruent; Incon = Incongruent. Error bars are one-tailed 95% confidence intervals. For individual patient data, see Figure 3 and Appendix.

The 2×2 ANOVA on Novel conditions showed a significant main effect of Congruency, F(1,28) = 21.2, p<.001, which reflected longer RTs in the Incongruent than Congruent condition, as expected (and paralleling the increased error rate reported above). There was also a significant main effect of Group, F(1,28) = 7.81, p<.01, which reflected longer RTs in Patients than Controls. There was no evidence for an interaction between Congruency and Group, F(1,28)<1. The slower RTs to Novel items for both Incongruent relative to Congruent conditions, and Patients relative to Controls, reinforces the importance of measuring priming by proportional as well as subtractive means below.

### Proportional Priming

The 2×2×2 ANOVA on proportional priming showed a significant main effect of Congruency, F(1,28)=23.2, p<.001, with positive priming (response speeding) for Congruent conditions and negative priming (response slowing) for Incongruent conditions, together with a significant main effect of Format Match, F(1,28)=30.5, p<.001, with more positive priming within formats rather than across formats, as expected. Unlike Horner and Henson (2011), any interaction between Congruency and Format Match did not reach significance, F(1,28)<1. There was a significant main effect of Group, F(1,28)=4.38, p<.05, with patients showing more negative priming scores on average (driven by Incongruent conditions; see Figure 2). Interestingly, there was no evidence for an interaction between Group and Congruency, F(1,28)=1.55, p=.224, nor for an interaction between Group and Format Match, F(1,28)=1.43, p=.241, nor for the three-way interaction, F(1,28)=1.24, p=.274. These results suggest that patients exhibit similar proportional priming effects as controls, i.e, were equally sensitive to S-R effects (see also Bayes Factor analyses below).

The significance of proportional priming effects in each of the 4 conditions separately is illustrated in Figure 2 (error bars are 95% confidence intervals, one-tailed, given the prior patterns in Horner & Henson, 2011). Like the young controls in Horner and Henson (2011), the present older controls showed significant positive priming in the two Congruent conditions, and significant negative priming in the Incongruent Across-format condition. The patients showed significant positive priming in the Congruent Within-format condition, but not the Congruent Across-format condition. The patients also showed negative priming, though this only reached significance in the Within-format Incongruent condition (p=.054 for the Across-format Incongruent condition). Thus while there was no evidence from the previous ANOVA that the pattern of priming differed across Groups, it is important that Patients did show cases of both positive and negative priming that were significant even when patients were considered on their own.

We conducted various additional analyses to confirm the significance of the proportional priming results.

### Subtractive Priming

We first analysed a subtractive rather than proportional measure of priming. The same 2×2×2 ANOVA on subtractive priming showed a significant main effect of Congruency, F(1,28)=5.84, p<.05, together with a significant main effect of Format Match, F(1,28)=13.2, p<.001, and highly significant main effect of Group, F(1,28)=9.98, p<.005. Again, any interaction between Congruency and Format Match did not reach significance; nor did the three-way interaction between Congruency, Format Match and Group, F(1,28)’s<1. The twoway interactions between Group and Congruency, F(1,28)=5.15, p<.05, and between Group and Format Match, F(1,28)=5.87, p<.05, did reach significance, unlike the results for proportional priming. Importantly though, these interactions reflected a larger effect of Congruency and of Format Match on subtractive priming for Patients than for Controls. In other words, subtractive priming suggested that Patients were actually *more* sensitive to S-R effects than were Controls (if slower RTs overall do not matter).

### Nonparametric Tests

Given that the patient group only had 6 members, while the control group had 24, it is difficult to assess the homogeneity of variance assumed by the above (parameteric) ANOVAs. We therefore performed non-parametric, Wilcoxon ranksum tests on proportional priming scores, separately for each of the within-group effects of 1) Congruency, 2) Format Match and 3) Congruency-by-Format-Match interaction. When combining both groups, there were significant main effects of Congruency, ranksum = 427, p < .001, and Format Match, ranksum = 437, p < .001, but no interaction between these two factors, ranksum = 281, p = .32. When comparing the two groups however, there was no evidence for any interaction between Group and any of these three effects, ranksums < 346, p’s > .19. The same pattern of significant results was obtained when using nonparametric analysis of subtractive priming scores.

### Bayes Factor

The lack of a significant two-way interaction between Congruency and Group suggests that the congruency effect, as an index of S-R bindings, is comparable in our patients compared to our controls. To provide more evidence for this conclusion, rather than relying on failure to reject the null hypothesis that the two groups differ, we calculated the Bayes Factor for the likelihood of patients showing the same size congruency effect as controls (alternate hypothesis), relative to the likelihood of patients showing no congruency effect (null hypothesis; Dienes, 2011). Given the congruency effect for proportional priming (averaged across format) had a mean of 19.4% and standard deviation of 21.4% in Controls, and a mean of 33.8% and standard deviation of 38.7% in Patients, the Bayes Factor was 5.09, which provides “substantial” (Jeffreys, 1961) evidence for the hypothesis that controls and patients show the same size congruency effect.

The results of the same analyses repeated with sex as a covariate were very similar (see Appendix).

### Comparison with Schnyer et al (2006)

For direct comparison with Experiment 2 of Schnyer et al (2006), we re-plotted the proportional priming scores from their study together with those from the present study. We took data from their Block 1, and from our Within-Format conditions, since these are the most comparable conditions. We averaged across the Low and High primed conditions of Schnyer et al (2006), given that these corresponded to 1 and 3 presentations, whereas we used 2 presentations. The results are shown in Figure 3. The main difference for the Controls is that the Incongruent condition abolished priming in the present study, but not the Schnyer et al study, consistent with our claim that the present design reverses multiple levels of response representation, including Stimulus-Classification bindings, which are not reversed in the Inverted condition of Schnyer et al. More importantly, reversing multiple levels of response representation also reduced priming in all six of the present Patients, unlike the patients in the Schnyer et al study, who showed no effect of decision and action reversal.

**Figure 3.**
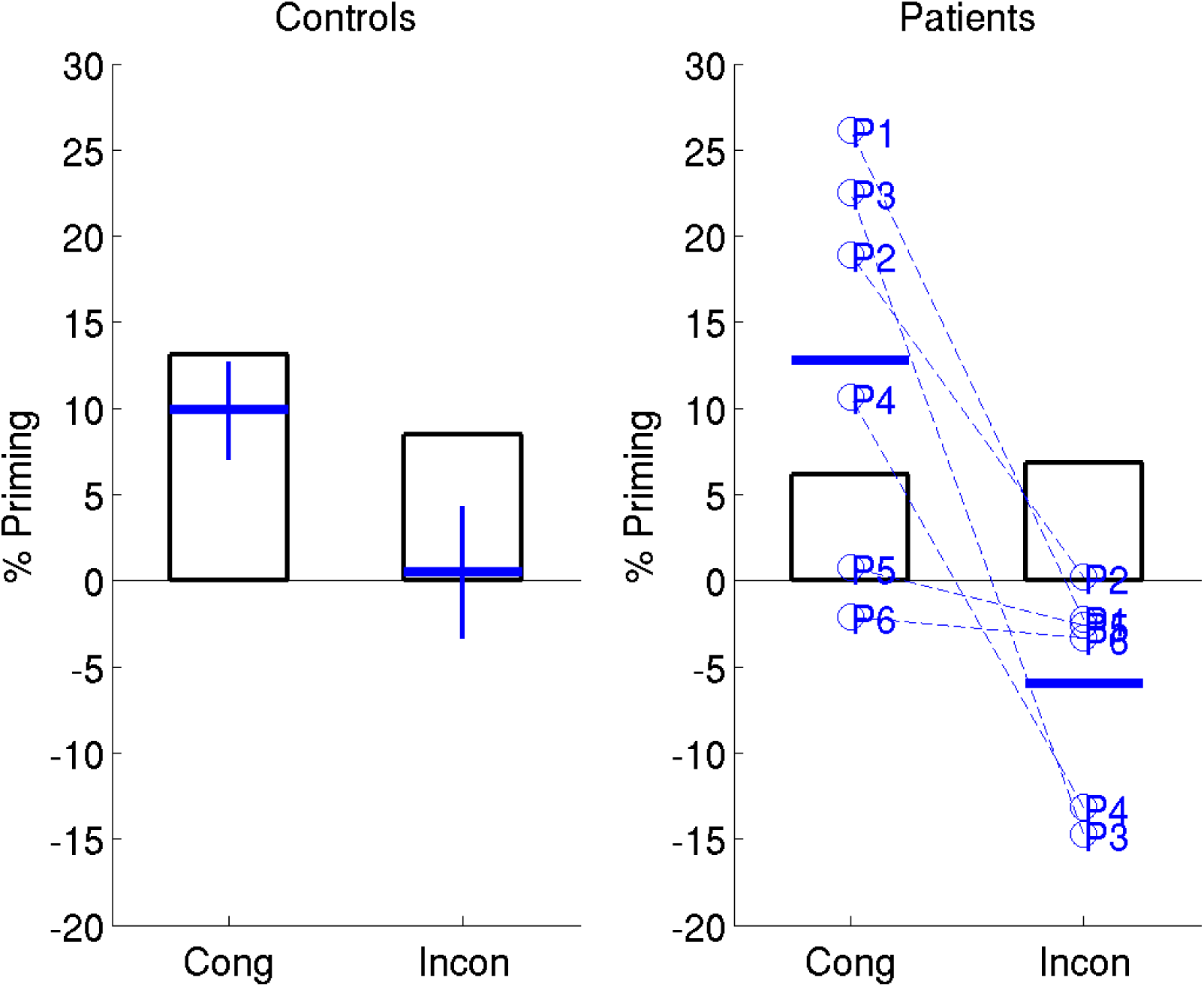
Comparison of proportional priming effects in current study with that of Schnyer et al. (2006; Experiment 2). The open bars are data replotted from Block 1 of Schnyer et al., averaging over their Low and High Primed conditions (N=12 Controls and N=9 Patients; no SD data provided); the blue horizontal lines and error bars are means and 95% one-tailed confidence intervals from the Within Format condition of the present study (N=24 Controls and N=6 patients), after adjusting for sex. Data from the six patients (P1-P6) from the present study are also plotted separately.

## Discussion

We hypothesized, based on Schnyer et al (2006), that individuals with hippocampal damage would show reduced evidence of stimulus-response (S-R) retrieval. S-R retrieval was indexed by the effect of congruency on RT priming, which reflects the difference in priming when the response is reversed between initial and repeated stimulus presentations, relative to when it is maintained. This hypothesis was not supported, in that we found that the evidence favoured the null hypothesis of equivalently-sized congruency effects in patients and controls, and the patients clearly showed significant congruency effects, no matter how we measured them. This finding is surprising, because according to many theories, the hippocampus would seem to be a critical brain structure for rapid learning (from just 1-2 presentations) of novel and arbitrary S-R bindings.

We start by considering possible reasons for the discrepancy between the Schnyer et al (2006) study and the present study, before discussing the broader implications of our findings.

### Possible differences from Schnyer et al. (2016)

One possible difference between the present study and that of Schnyer et al (2006) concerns the type of patients. For example, the patients tested by Schnyer et al (2006) may have had more severe or extensive lesions of hippocampus, such that a threshold for observing reduced S-R bindings was exceeded. A direct comparison is not possible because Schnyer et al (2006) did not report details of the extent of hippocampal damage. However it is noteworthy that the average, bilateral gray-matter volume loss in hippocampus was 40% in the present cases, which is comparable to other studies of acquired hippocampal damage in human adults (Henson et al, 2016). Moreover, all six of the present cases showed marked impairment on at least one standardised neuropsychological test of memory (though unfortunately, these were different tests from those used by Schnyer et al., so again, direct comparison is difficult). Thus while we cannot refute the possibility that the patients in the Schnyer et al study had greater hippocampal damage and more severe amnesia, the present hippocampal lesions were sufficiently severe to impair episodic memory, and so it remains interesting that there was no corresponding impairment detected on S-R binding.

Another possibility is that the extent of damage outside the hippocampus was greater in Schnyer et al’s patients; for example, spreading into surrounding rhinal cortex. This would be consistent with lesion work in non-human primates that has investigated rapid learning of arbitrary visuo-motor mappings (visuomotor learning, VML), which resemble S-R bindings. Although initial work using aspiration lesions of the hippocampus plus underlying cortex, including caudal entorhinal cortex, indicated that these regions were critical (Murray and Wise, 1996; Murray and Mishkin, 1998), subsequent research using more precise, reversible inactivation via intracerebral infusion of a GABA agonist found that hippocampus was not essential for VML; rather entorhinal cortex was key (Yang *et al.*, 2014). Only 2 of the 6 patients in the present study (P4 and P6) had significant entorhinal volume loss detectable from MRI (compared to all 6 for hippocampal volume loss, Henson et al, 2016). It is also possible that perirhinal cortex plays a role in binding stimuli and responses, given evidence that this region supports not only representations of complex visual objects, but also the encoding of object-context associations (Watson *et al.*, 2012). If the patients in Schnyer et al.’s study had greater ento- or peri-rhinal damage, then they would show stronger evidence of impairment of S-R bindings. Note that, while this would reconcile results across the two studies, the present study would still be correct in concluding that hippocampus proper is not necessary for S-R bindings.

Other NHP studies have shown that the fornix is also critical for VML (Brasted *et al.*, 2005), yet diffusion-weighted MRI showed that the present group of 6 patients also had significant white matter abnormalities of the fornix, in addition to their hippocampal damage (Henson et al, 2016). Thus the relationship between the neural correlates of S-R bindings in humans and neural correlates of VML in non-human primates requires further investigation.

A final anatomical possibility is that the patients in Schnyer et al.’s study (but not those in the present study) also had damage to basal ganglia circuits that support many types of motor/habit learning (e.g., Gabrieli, 1998; Poldrack *et al.*, 2001). However, motor (procedural) learning is conventionally thought to be more gradual (i.e., incremental over many trials) than the type of S-R learning occurring here; a point we return to later.

A second dimension along which the two studies differ concerns the paradigm. We reversed response contingencies by changing the referent of the size judgment task, rather than reversing the task instruction. As explained in the Introduction, the reason for this change was that we have previously shown (Horner & Henson, 2009, 2011) that a referent change additionally invalidates any bindings between the stimulus and the “classification” response (i.e., the label “bigger” or “smaller”), so the present referent change arguably induces a more comprehensive disruption of the use of S-R bindings. This could explain why priming was completely abolished in the present Incongruent (Within Format) condition, whereas it remained significant (though reduced) in the corresponding (Inverted) condition of the Schnyer et al. (2006) study (indeed, the use of Stimulus-Classification, S-C, bindings might be the cause of this residual priming in Schnyer et al.’s data, rather than a residual perceptual processing component). However, if the hippocampus is necessary specifically for S-C bindings, this would predict the opposite pattern of results across the two studies: ie, reduced congruency effects in present study, but equivalent congruent effects in Schnyer et al. study. Nonetheless, it remains possible that reversing the task instructions has a qualitatively different effect (e.g, in terms of task switching, Henson et al., 2016) from changing the referent; more specific hypotheses would be needed to test this.

Another difference between the two paradigms concerns the number of study presentations (before the task or referent is changed). Schnyer et al. used both a single presentation (their Low Prime condition) and three presentations (their High Prime condition), whereas we used two presentations. Thus one possibility is that two study repetitions were sufficient for non-hippocampal S-R learning (e.g., via basal ganglia). However, this is contrary to the effect that the number of presentations had on Schnyer et al’s results: Although they did not analyse their High and Low conditions separately, the numerical pattern for their single presentation condition actually showed a small cost of task reversal in Block 1 for both patients and controls; it was only after three presentations that Schnyer et al observed a greater task reversal cost in controls than patients. Thus the data from our study appear more like those from Schnyer et al’s Low Prime condition, rather than their High Prime condition. Another possible way to reconcile the two studies therefore is to propose that the difference between patients and controls only emerges after three or more presentations. In other words, three presentations may be necessary for controls to form an explicit representation of an S-R contingency in the hippocampus. However, this is the opposite to 1) conventional views that the hippocampus supports one-shot learning, and that other (eg basal ganglia) systems support more gradual, procedural learning (Gabrieli, 1998), and 2) other studies in healthy controls, which found only quantitative rather than qualitative effects of a small number of repetitions on S-R learning (e.g., Horner & Henson, 2009; Dennis & Perfect, 2012).

A final possibility is that the discrepant results across the two studies reflect statistical artefacts. It is possible that our failure to find an interaction between controls vs patients and congruent vs incongruent conditions was a type II error (given that we only tested 6 patients, whereas Schnyer et al tested 9). However, our Bayes Factor analysis favours the null hypothesis of no interaction relative to the alternative of an interaction the size that Schnyer et al found. Moreover, the three-way interaction between Group, Cue (Congruency) and Condition (Number of Repetitions) was not actually significant in Schnyer et al’s study either; their main claim was based on the fact that their Control group showed a significant Cue-by-Condition interaction, but their Patient group did not. Thus it is also possible that Schnyer et al’s failure to find evidence for S-R bindings in the patients was a type II error (whereas here, all 6 of the patients showed evidence of S-R bindings).

### Implications

We think that evidence for S-R bindings, like that presented here, implies an impressive ability of the human brain to rapidly learn a large number of S-R mappings. Of course, we do not know that an S-R binding was formed or retrieved for every trial in the experiment – it is possible that only a small fraction of trials in which S-R retrieval occurred was sufficient to cause the average RT priming effects – but at the other extreme, the implication is that the brain can store hundreds of unique mappings. So if the hippocampus is not the brain structure that supports this impressive feat, which brain region is?

As mentioned above, one possibility is the surrounding entorhinal or perirhinal cortex, consistent with the animal lesions studies using the VML task, while another possibility are basal ganglia structures, which have previously been associated with procedural learning (though normally associated with more gradual learning, they may be able to learn enough from 1-2 stimulus-response pairings to produce a detectable effect on RTs). A third possibility, not considered above, is that S-R bindings are mediated by prefrontal regions. This would be consistent with human fMRI and M/EEG studies of S-R retrieval, which implicate ventral prefrontal regions in particular (e.g., Horner & Henson, 2008; 2012; Race *et al.*, 2009; Wig *et al.*, 2009). It would also be consistent with evidence that ventral (and orbital) prefrontal lesions in animals impair VML (Bussey *et al.*, 2001). It would therefore be interesting to run the present paradigm in human patients with prefrontal lesions, where significantly smaller congruency effects would be predicted.

Theoretically, S-R bindings may not conform to the type of flexible associations attributed to hippocampus (Cohen *et al.*, 1997); associations that can be voluntarily retrieved and inter-related (e.g, to make transitive inferences across associations). While S-R bindings are clearly complex, encoding several types of response representations (Horner & Henson, 2009) and relatively abstract stimulus representations (as shown by the congruency effect in the present Across Format conditions), they may be relatively inflexible, in the sense of being retrieved automatically and independently. A related possibility is that S-R bindings are not represented explicitly, in the sense of participants being aware of them, which is why hippocampus is not involved. While we cannot rule out the possibility that both controls and patients had episodic memories for some trials (those in which S-R bindings were formed), the fact that S-R bindings co-occur with impairments on standard tests of episodic (explicit) memory in the present patients suggests that S-R bindings are more likely to be implicit (see also Giesen and Rothermund, 2015). In short, the inflexible, involuntary and/or implicit nature of S-R bindings may indicate a non-hippocampal locus; but nonetheless a locus that allows rapid encoding of multiple arbitrary mappings. This reinforces the point that it is theoretically important to understand not only what individuals with amnesia cannot to, but also what they can do (Clark and Maguire, 2016).

## Acknowledgements & Funding

This work was supported by the UK Medical Research Council (MC_A060_5PR10). We are very grateful to all the patients and their families for enabling this research. We also thank Dr Elisabeth Murray for helpful comments, and Dr Elias Mouchlianitis for helping with data collection. A.J.H. was supported in part by the Wellcome Trust (204277/Z/16/Z). J.S.S. was supported in part by a Scholar Award from the James S. McDonnell Foundation and in part by BBSRC grant number BB/L02263X/1. The full data and analysis scripts will be made available on publication.

## Appendix

**Table 1:** Each patient’s error and RT data separately

| Condition / | Within-format<br>(Picture-Picture) |  | Across-format<br>(Word-Picture) |  | Novel |  |
| --- | --- | --- | --- | --- | --- | --- |
| Congruency | Con | Incon | Con | Incon | Con | Incon |
| *% Errors* |  |  |  |  |  |  |
| *P1* | 2 | 17 | 3 | 11 | 6 | 8 |
| *P2* | 3 | 13 | 6 | 6 | 0 | 9 |
| *P3* | 8 | 30 | 3 | 19 | 2 | 20 |
| *P4* | 5 | 14 | 6 | 14 | 2 | 22 |
| *P5* | 5 | 8 | 8 | 11 | 9 | 13 |
| *P6* | 3 | 22 | 2 | 25 | 6 | 23 |
| *RTs (ms)* |  |  |  |  |  |  |
| *P1* | 783 | 1155 | 974 | 1000 | 1042 | 1086 |
| *P2* | 1146 | 1834 | 1549 | 2127 | 1410 | 1790 |
| *P3* | 3810 | 5329 | 4032 | 6893 | 4906 | 4623 |
| *P4* | 1066 | 1920 | 1313 | 2017 | 1204 | 1672 |
| *P5* | 1029 | 1241 | 1114 | 1313 | 1009 | 1259 |
| *P6* | 765 | 977 | 836 | 978 | 777 | 904 |

Given that one of the six patients were female, yet 13 of the 24 controls were female, the main analyses were repeated with sex as a covariate.

### Error Analyses

The 2×2×2 ANOVA on subtractive priming of errors showed no significant effects or interactions between Format Match, Congruency and Group, F(1,27)’s < 1.87, p’s > .18, matching the pattern without adjusting for sex in the main paper.

### RT Analyses

The 2×2×2 ANOVA on proportional priming showed a significant main effect of Congruency, F(1,27)=13.7, p<.001, with positive priming (response speeding) for Congruent conditions and negative priming (response slowing) for Incongruent conditions, together with a significant main effect of Format Match, F(1,27)=23.1, p<.001, with more positive priming within formats rather than across formats, as expected. The interaction between Congruency and Format Match did not reach significance, F(1,27)<1. The main effect of Participant Group was not significant, F(1,27)=2.24, p=.146, and nor was there evidence for an interaction between Participant Group and Congruency, F(1,27)<1, nor between Participant Group and Format Match, F(1,27)<1, nor three-way interaction, F(1,27)=1.19, p=.177. Thus the only change after covarying out sex was that the main effect of participant group was no longer significant, but this only bolsters the general claim of the present study that patients perform similarly to controls.

For subtractive priming, the same 2×2×2 ANOVA on subtractive priming showed a significant main effect of Congruency, F(1,27)=5.81, p<.05, together with a significant main effect of Format Match, F(1,27)=13.0, p<.001, and significant main effect of Group, F(1,27)=6.52, p<.05. Again, any interaction between Congruency and Format Match did not reach significance; nor did the three-way interaction between Congruency, Format Match and Group, F(1,27)’s<1. The two-way interactions between Group and Congruency, F(1,27)=3.71, p=.065, and between Group and Format Match, F(1,27)=4.53, p<.05, approached or reached significance, unlike the results for proportional priming, but like for the analysis of subtractive priming without adjustment for sex. Again though, these interactions reflected a larger effect of Congruency and of Format Match on subtractive priming for Patients than for Controls, i.e, Patients were actually *more* sensitive to S-R effects than were Controls (if slower RTs overall do not matter).

## References

Brasted PJ, Bussey TJ, Murray EA, Wise SP. Conditional motor learning in the nonspatial domain: effects of errorless learning and the contribution of the fornix to one-trial learning [Internet]. Behav. Neurosci. 2005; 119: 662–676. Available from: papers2://publication/doi/10.1037/0735-7044.119.3.662

Bussey TJ, Wise SP, Murray E a. The role of ventral and orbital prefrontal cortex in conditional visuomotor learning and strategy use in rhesus monkeys (Macaca mulatta). Behav. Neurosci. 2001; 115: 971–982.

Cave CB, Squire LR. Intact and long-lasting repetition priming in amnesia. J. Exp. Psychol. Learn. Mem. Cogn. 1992; 18: 509–520.

Clark I a., Maguire E a. Remembering Preservation in Hippocampal Amnesia [Internet]. Annu. Rev. Psychol. 2016; 67: annurev-psych-122414-033739. Available from: http://www.annualreviews.org/doi/10.1146/annurev-psych-122414-033739

Cohen NJ, Eichenbaum HB. Memory, amnesia and the hippocampal system. MIT Press 1994: 55173–55173.

Cohen NJ, Poldrack R a., Eichenbaum H. Memory for Items and Memory for Relations in the Procedural/Declarative Memory Framework [Internet]. Memory 1997; 5: 131–178.Available from: http://dx.doi.org/10.1080/741941149%5Cnhttp://www.tandfonline.com/doi/abs/10.1080/741941149%5Cnhttp://www.tandfonline.com/doi/pdf/10.1080/741941149

Denkinger B, Koutstaal W. Perceive-decide-act, perceive-decide-act: how abstract is repetition-related decision learning? J. Exp. Psychol. Learn. Mem. Cogn. 2009; 35: 742–756 .

Dennis I, Perfect TJ. Do Stimulus-Action Associations Contribute to Repetition Priming? J. Exp. Psychol. Learn. Mem. Cogn. 2012; 39: 85–95 .

Dobbins IG Schnyer DM, Verfaellie M, Schacter DL. Cortical activity reductions during repetition priming can result from rapid response learning. [Internet]. Nature 2004; 428: 316–9.Available from: http://www.ncbi.nlm.nih.gov/pubmed/14990968

Gabrieli JD. Cognitive neuroscience of human memory. Annu. Rev. Psychol. 1998; 49: 87–115.

Giesen C, Rothermund K. Adapting to stimulus-response contingencies without noticing them. [Internet]. J. Exp. Psychol. Hum. Percept. Perform. 2015; 41: 1475–1481.Available from: http://doi.apa.org/getdoi.cfm?doi=10.1037/xhp0000122

Henson RNA. Neuroimaging studies of priming. Prog. Neurobiol. 2003; 70: 53–81.

Henson RN, Eckstein D, Waszak F, Frings C, Horner AJ. Stimulus-response bindings in priming [Internet]. Trends Cogn. Sci. 2014; 18: 376–383.Available from: http://dx.doi.org/10.1016/_j.tics.2014.03.004

Henson RN, Greve A, Cooper E, Gregori M, Simons JS, Geerligs L, et al. The effects of hippocampal lesions on MRI measures of structural and functional connectivity [Internet]. Hippocampus 2016; 1463: 1–48.Available from: http://doi.wiley.com/10.1002/hipo.22621

Horner AJ, Henson RN. Priming, response learning and repetition suppression. Neuropsychologia 2008; 46: 1979–1991.

Horner AJ, Henson RN. Bindings between stimuli and multiple response codes dominate long-lag repetition priming in speeded classification tasks. J. Exp. Psychol. Learn. Mem. Cogn. 2009; 35: 757–779.

Horner AJ, Henson RN. Stimulus-response bindings code both abstract and specific representations of stimuli: evidence from a classification priming design that reverses multiple levels of response representation. Mem. Cognit. 2011; 39: 1457–1471: .

Horner AJ, Henson RN. Incongruent abstract stimulus-response bindings result in response interference: fMRI and EEG evidence from visual object classification priming. J. Cogn. Neurosci. 2012; 24: 760–773.

Koutstaal W, Wagner AD, Rotte M, Maril A, Buckner RL, Schacter DL. Perceptual specificity in visual object priming: Functional magnetic resonance imaging evidence for a laterality difference in fusiform cortex. Neuropsychologia 2001; 39: 184–199.

Murray EA, Mishkin M. Object Recognition and Location Memory in Monkeys with Excitotoxic Lesions of the Amygdala and Hippocampus. 1998; 18: 6568–6582.

Murray E, Wise S. Role of the hippocampus plus subjacent cortex but not amygdala in visuomotor conditional learning in rhesus monkeys. [Internet]. Behav. Neurosci. 1996; 110: 1261–1270:.Available from: http://psycnet.apa.org/psycinfo/1996-07070-005%5Cnpapers2://publication/uuid/AFE8FF99-F7D3-4478-A194-D5C395FA4115

Poldrack RA, Clark J, Paré-Blagoev EJ, Shohamy D, Creso Moyano J, Myers C, et al. Interactive memory systems in the human brain. [Internet]. Nature 2001; 414: 546–550.Available from: http://www.ncbi.nlm.nih.gov/sites/entrez?Db=pubmed&DbFrom=pubmed&Cmd=Link&LinkName=pubmed_pubmed&LinkReadableName=RelatedArticles&IdsFromResult=11734855&ordinalpos=3&itool=EntrezSystem2.PEntrez.Pubmed.Pubmed_ResultsPanel.Pubmed_RVDocSum

Race E a, Shanker S, Wagner AD. Neural priming in human frontal cortex: multiple forms of learning reduce demands on the prefrontal executive system. J. Cogn. Neurosci. 2009; 21: 1766–1781.

Schacter DL, Buckner RL. Priming and the brain. Neuron 1998; 20: 185–195.

Schacter DL, Chiu CY, Ochsner KN. Implicit memory: a selective review. Annu. Rev. Neurosci. 1993; 16: 159–182.

Schacter DL, Tulving E. What are the memory systems of 1994? [Internet]. Mem. Syst. 1994. 1994: 1–38.Available from: http://search.ebscohost.com/login.aspx?direct=true&db=psyh&AN=1994-98504-001&site=ehost-live

Schnyer DM, Dobbins IG Nicholls L, Davis S, Verfaellie M, Schacter DL. Item to decision mapping in rapid response learning. Mem. Cognit. 2007; 35: 1472–1482.

Schnyer DM, Dobbins IG Nicholls L, Schacter DL, Verfaellie M. Rapid response learning in amnesia: Delineating associative learning components in repetition priming. Neuropsychologia 2006; 44: 140–149.

Simons JS, Koutstaal W, Prince S, Wagner AD, Schacter DL. Neural mechanisms of visual object priming: Evidence for perceptual and semantic distinctions in fusiform cortex. Neuroimage 2003; 19: 613–626.

Squire LR. Memory and the hippocampus: a synthesis from findings with rats, monkeys, and humans. Psychol. Rev. 1992; 99: 195–231.

Watson HC, Wilding EL, Graham KS. A Role for Perirhinal Cortex in Memory for Novel Object-Context Associations [Internet]. J. Neurosci. 2012; 32: 4473–4481.Available from: http://www.jneurosci.org/cgi/doi/10.1523/JNEUROSCI.5751-11.2012

Wig GS, Buckner RL, Schacter DL. Repetition priming influences distinct brain systems: evidence from task-evoked data and resting-state correlations. J. Neurophysiol. 2009; 101: 2632–48.

Yang T, Bavley RL, Fomalont K, Blomstrom KJ, Mitz AR, Turchi J, et al. Contributions of the hippocampus and entorhinal cortex to rapid visuomotor learning in rhesus monkeys. [Internet]. Hippocampus 2014; 24: 1102–11.[cited 2014 Sep 28] Available from: http://www.ncbi.nlm.nih.gov/pubmed/24753214

